# Signal Transduction in an Enzymatic Photoreceptor Revealed by Cryo-Electron Microscopy

**DOI:** 10.1101/2023.11.08.566274

**Authors:** Tek Narsingh Malla, Carolina Hernandez, David Menendez, Dorina Bizhga, Joshua H. Mendez, Srinivasan Muniyappan, Peter Schwander, Emina A. Stojković, Marius Schmidt

## Abstract

Phytochromes are essential photoreceptor proteins in plants with homologs in bacteria and fungi that regulate a variety of important environmental responses. They display a reversible photocycle between two distinct states, the red-light absorbing Pr and the far-red light absorbing Pfr, each with its own structure. The reversible Pr to Pfr photoconversion requires covalently bound bilin chromophore and regulates the activity of a C-terminal enzymatic domain, which is usually a histidine kinase (HK). In plants, phytochromes translocate to nucleus where the C-terminal effector domain interacts with protein interaction factors (PIFs) to induce gene expression. In bacteria, the HK phosphorylates a response-regulator (RR) protein triggering downstream gene expression through a two-component signaling pathway. Although plant and bacterial phytochromes share similar structural composition, they have contrasting activity in the presence of light with most BphPs being active in the dark. The molecular mechanism that explains bacterial and plant phytochrome signaling has not been well understood due to limited structures of full-length phytochromes with enzymatic domain resolved at or near atomic resolution in both Pr and Pfr states. Here, we report the first Cryo-EM structures of a wild-type bacterial phytochrome with a HK enzymatic domain, determined in both Pr and Pfr states, between 3.75 and 4.13 Å resolution, respectively. Furthermore, we capture a distinct Pr/Pfr heterodimer of the same protein as potential signal transduction intermediate at 3.75 Å resolution. Our three Cryo-EM structures of the distinct signaling states of BphPs are further reinforced by Cryo-EM structures of the truncated PCM of the same protein determined for the Pr/Pfr heterodimer as well as Pfr state. These structures provide insight into the different light-signaling mechanisms that could explain how bacteria and plants see the light.

## 1. Introduction

The ability to sense and respond to changing environmental conditions is essential to the survival of organisms. In bacteria, environmental signals are typically sensed through a two-component signaling mechanism composed of a histidine kinase (HK) and a response regulator (RR) (Stock, Robinson et al. 2000). Two-component systems are highly modular and have been adapted into a variety of cellular signaling circuits. The majority of enzymatic photoreceptors in bacteria are part of such a signaling mechanism by adapting to changes in the spectrum, the intensity, and the direction of light (Furuya 1993). Phytochromes are ubiquitous among plants and widespread in fungi and bacteria (Davis, Vener et al. 1999, Rockwell, Su et al. 2006). The plant phytochromes (PHYs) have a different composition (Fig. 1 a,b) and may also display different structures (Li, Burgie et al. 2022). Upon light activation plant PHYs migrate to the nucleus where they induce responses such as germination, greening and shade avoiding (Batschauer 1998) in a mechanism distinct from the bacterial two-component system. In photosynthetic bacteria, phytochromes stimulate the synthesis of light-harvesting complexes (Giraud, Fardoux et al. 2002, Giraud, Zappa et al. 2005). In non-photosynthetic bacteria their role is not as well understood (Wagner, Brunzelle et al. 2005, Woitowich, Halavaty et al. 2018). Here, details of the signaling mechanism in the nonphotosynthetic myxobacterium *Stigmatella aurantiaca* (Sanchez, Carrillo et al. 2019) are presented. Myxobacteria are distinguished among prokaryotes by a multicellular stage in their life cycle known as fruiting bodies. In *S. aurantica* fruiting body formation is controlled by red and far-red light, implicating bacteriophytochrome (BphP) signaling (Qualls, Stephens et al. 1978, White, Shropshire et al. 1980, Woitowich, Halavaty et al. 2018). Myxobacteria are also extensively studied for the synthesis of secondary metabolites with anticancer and antimicrobial properties, recently discovered to be modified by light (Schaberle, Lohr et al. 2014, Lapuhs, Heinrich et al. 2022).

**Figure 1.**
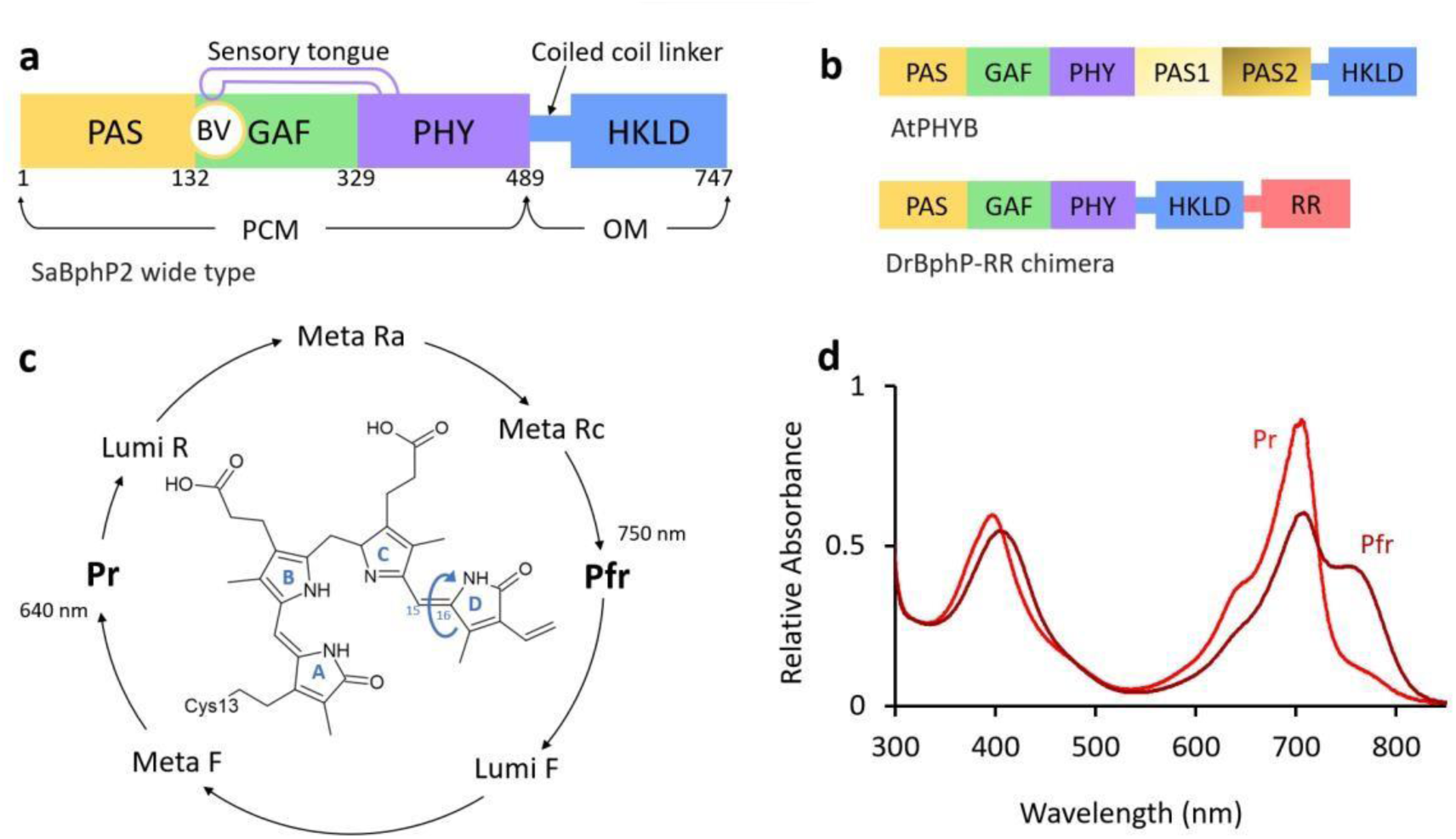
*S. aurantiaca* BphP2 domain composition and Pr/Pfr photocycle. (a) Schematic presentation of domain architecture of the wild -type SaBphP2. Sequence numbers are marked for each domain. PAS, GAF and PHY domains form the PCM which is covalently linked to the output effector module (OM). In SaBphP2, the OM is a HK like domain (HKLD). The BV chromophore is covalently bound to the conserved cysteine N-terminal of the PAS domain. The BV binding pocket is within the GAF domain. The PHY domain has a long hair-pin extension called the sensory tongue that is required for complet Pr/Pfr photoconversion. (b) Domain composition of the *Arabidopsis thaliana* plant PHYB and the *Deinococcus radiodurans* BphP-RR. chimera; (c) SaBphP2 reversible photocycle with respective intermediates. When the Pr state is illuminated with red light (640 nm - 700 nm), it converts to the Pfr state. When illuminated with far -red light (750 nm), the Pfr state can be driven to Pr. The structure of BV is shown with the 4 pyrrole rings. The C _15_=C_16_ double bond between rings C and D that undergoes Z to E isomerization is marked by an arrow. (d) Absorption spectra of SaBphP2 with red (700 nm) and far-red (750 nm) absorbing Pr and Pfr states, respectively.

BphPs consists of a photosensory core module (PCM) and an output effector module (OM) (Fig. 1 a). The PCM is composed of three domains: PAS (Per-Arndt-Sim), GAF (cGMP phosphodiesterase/adenylyl cyclase/FhIA), and PHY (phytochrome-specific GAF-related). The central chromophore, biliverdin (BV) (Fig. 1 c), is a heme-derived, open-chain tetrapyrrole (pyrrole rings A–D) that is covalently bound to an Nterminal conserved cysteine. The C-terminal OM is covalently bound to the PHY domain and differs among species, although a HK domain is the most common (Rockwell, Su et al. 2006, Kojadinovic, Laugraud et al. 2008, Otero, Klinke et al. 2016). In *Stigmatella aurantica* two BphPs with similar domain composition are found (Fig. 1 a). In the classical bacteriophytochrome 2 (SaBphP2), the C-terminal OM is a putative HK denoted HK-like domain (HKLD) in the following. SaBphP2 is part of a single operon containing downstream genes encoding a RR for the two-component signaling pathway and a heme oxygenase required for BV synthesis (Woitowich, Halavaty et al. 2018).

BphPs interconvert between a red-light absorbing, Pr state and a far red-light absorbing Pfr state, through several intermediates (Fig. 1 c,d) (Auldridge and Forest 2011). Upon illumination with red light (λ = 640 nm), the BV undergoes a Z to E isomerization about the C_15_=C_16_ double bond between the pyrrole rings C and D (Fig. 1 c) driving the phytochrome into the first half of the photocycle (Takala, Bjorling et al. 2014, Burgie, Zhang et al. 2016, Carrillo, Pandey et al. 2021). Following successive deprotonation and reprotonation events of BV in the Meta states (Auldridge and Forest 2011), rearrangement of the water network and the conserved amino acids in the BV-binding pocket and the neighboring sensory tongue of the PHY domain, the phytochrome reaches the Pfr state (Carrillo, Pandey et al. 2021) ultimately controlling the activity of the C-terminal OM. The sensory tongue of the PHY domain (Fig. 1 a) shifts between a βstrand conformation in the Pr and an α-helical conformation in the Pfr state as demonstrated by several structural and spectroscopic studies (Takala, Bjorling et al. 2014, Bjorling, Berntsson et al. 2015, Burgie, Zhang et al. 2016, Wahlgren, Claesson et al. 2022). A sequence of amino acid residues called the PRXSF motif in the PHY domain (Essen, Mailliet et al. 2008, Stojkovic, Toh et al. 2014) plays an important role in this transition, and henceforth in the sensing of light. The Arg of the PRXSF motif forms a key salt bridge with Asp of the PASDIP motif (Yang, Stojkovic et al. 2007) of the BV-binding GAF domain in the darkadapted Pr state. This Arg-Asp salt bridge is broken upon the β-strand to α-helix conformational change in the Pfr state (Fig. 1 b). In the dark, classical BphPs like SaBphP2 undergo thermal reversion to the Pr-state.

## 2. Results

A comprehensive understanding of the signal transduction mechanism from the BV-binding PCM to the OM with enzymatic activity covering nearly 140 Å in distance has been limited by the structures at atomic and near-atomic resolution of the intact, wild-type BphPs in both, Pr and Pfr states. Here, we present distinct BphP structures and corresponding cryo-EM maps of the full-length canonical SaBphP2 containing a HKLD, determined at near atomic resolution using single-particle cryo-EM. The cryo-EM structures reveal conformational changes between different signaling states of the reversible phytochrome photocycle and provide concrete evidence for the highly coveted mechanism of light-induced signal transduction. Details of cryo-EM map reconstruction and structure determination are listed in the Supplementary Material.

### 2.1 The structure of the Pr homodimer

The structure of the full length SaBphP2 homodimer in the Pr state is shown in Fig. 2 a together with the corresponding cryo-EM map. The PCM portion of the map as well as part of the linker (up to residue 538) was interpreted with a structural model (green ribbons). The structure of the PCM part is similar to the published high-resolution crystal structure (Sanchez, Carrillo et al. 2019) as shown by the orange ribbon overlaid on subunit A (Fig. 2 a). The dimer interface of both PCM and the linker region to the HKLD domain, is composed of two long α-helices, one from each subunit.

**Figure 2.**
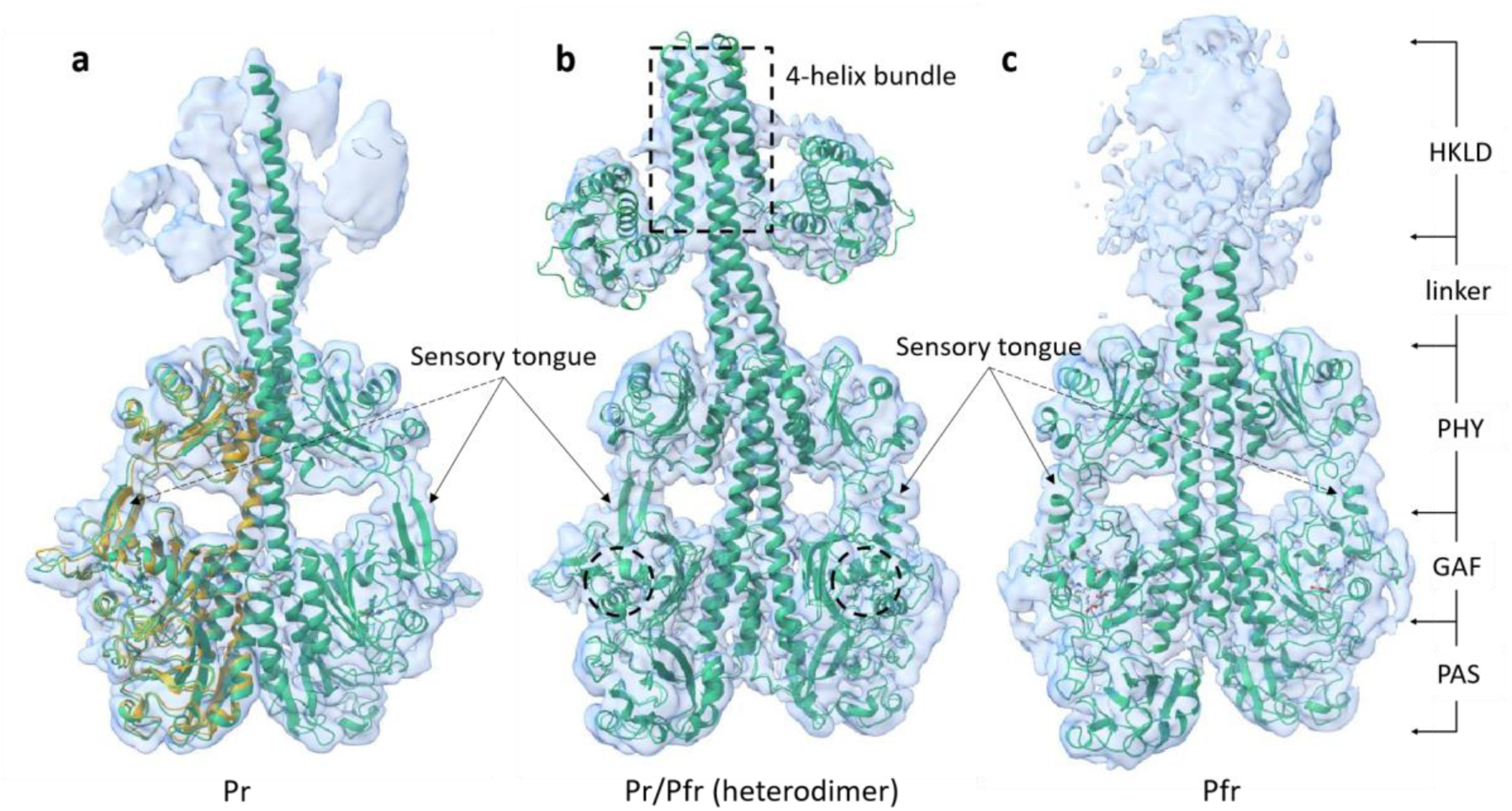
The cryo-EM-structures of full-length SaBphP2 dimer. The protein structures are shown by green ribbons with corresponding maps displayed in light blue, (a) In the Pr homodimer, the sensory tongues of the PHY domains in both subunits are in the β-sheet conformation. The monomer from the crystal structure of the SaBphP2 PCM dimer (PDB code: 6PTQ) in the Pr state (orange cartoon) is overlaid on subunit A. (b) In the SaBphP2-Pr/Pfr heterodimer, the configuration of the sensory tongue is an a-helix in one monomer and a β-strand in the other. The HKLD is well resolved. The 4-helix bundle is marked by a dashed box, the locations of the BV chromophore by dashed circles, (c) In the Pfr state homodimer, both sensory tongues are in the a-helix configuration. The approximate locations of the domains in the SaBphP2 dimer are marked on the right.

The sensory tongue of the PHY domain, in both monomers, is a two stranded β-sheet which is also observed in the crystal structure of SaBphP2 PCM in the Pr state. The length of the sensory tongue, measured by the distance between the top of the GAF domain (His187) and the bottom of the PHY domain (Thr470), is ∼18 Å (Fig. 3 d). The BV chromophore is in the all-Z (ssa) configuration with ring D forming hydrogen bond with the conserved His275 of the GAF domain (Fig. 3 a). The BV is further stabilized by additional conserved residues also shown in Fig. 3 a. The conserved Asp192 and His 245 are forming H-bonds with pyrrole rings A and C, respectively. The propionate of ring B is connected by hydrogen bonds to the conserved Tyr201 and Arg239, and that of ring C to Arg207, Ser257 and Ser259. The conserved pyrrole water near pyrrole rings A, B and C which has been observed in the crystal structures of the PCM of SaBphP2 and related BphPs (Edlund, Takala et al. 2016, Sanchez, Carrillo et al. 2019) was not resolved in the map. The conserved Arg457 of the PRXSF motif of the PHY sensory tongue probes the BV binding pocket by forming a salt bridge with GAF-domain Asp192.

**Figure 3.**
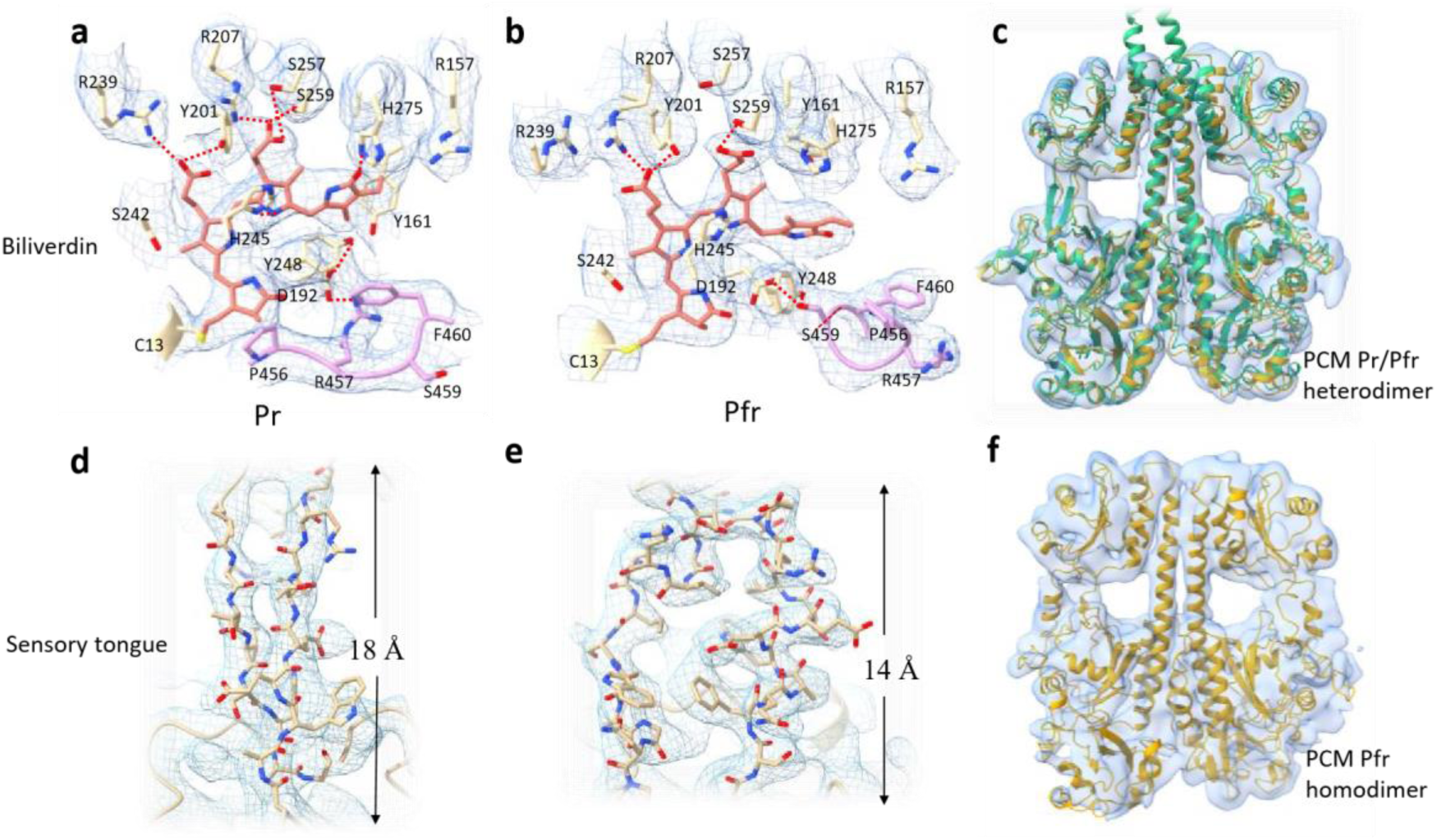
Pr /Pfr photoconversion in SaBphP2. The structures of the BV chromophore and the sensory tongue are shown with the respective CryoEM maps in light blue in (a) - (e). (a) The BV chromophore in the Pr state (orange). Conserved amino acid residues are marked. Ring D is in the Z configuration. The PRXSF motif with individual amino acid side chains is shown in pink. Hydrogen bonds are marked with red dotted lines, (b) The BV chromophore in the Pfr state (orange). Conserved amino acid residues are marked. Hydrogen bonds are shown with red dotted lines. The ring D is in the Z configuration. The PRXSF motif of the sensory tongue is shown in pink, (c) The PCM in the Pr/Pfr heterodimeric state (orange) is overlayed on the full-length SaBphP2 in the Pr/Pfr heterodimer state (green). The structures of full-length SaBphP2 and that of the PCM are essentially identical in the PCM region, (d) The sensory tongue structure is characterized by two β-strands in the Pr state, (e) In the Pfr state, the strands move apart and one of the strands forms an a-helix. The length of the sensory tongue is marked in (d) and (e). (f) The cryo-EM structure of the PCM only in the Pfr state.

To compare the structures, a minimalistic impression of the SaBphP2 architecture can be obtained by displaying the dimer interface helices which are shown in Fig. 4 in dark green (subunit A) and pink (subunit B) and the linker helices to the effector domain which are shown by a lighter shade of green and pink, respectively. In the Pr homodimer, the dimer and the linker helices overlap within the PHY domain. The top positions of the dimer interface helices are ∼25 Å apart (Fig. 4 a,g). The PAS and PHY domains are separated by 39 Å and 22 Å, respectively (Fig. 4 a,d). Fig. 5 provides a different view on the SaBphP2 structural building blocks. The four structural domains of the BphP are not lying all in the same plane. When measured with respect to the PAS domain, the GAF domain is oriented at an angle of 55˚, the PHY domain at 60˚ and the HKLD effector module at 35˚ (Fig. 5 a,d).

**Figure 4.**
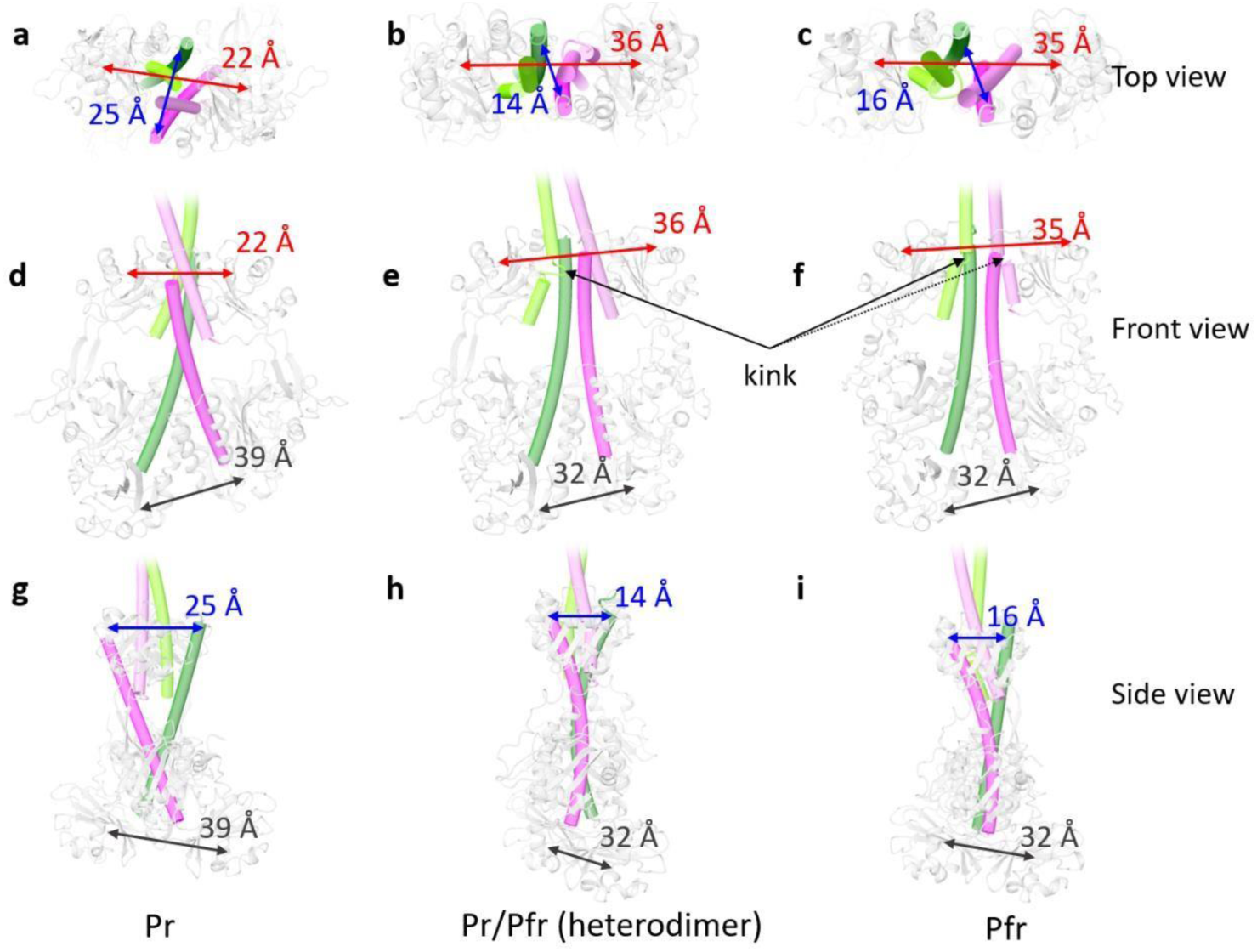
The dimer interface of the intact SaBphP2. The top (a-c), front (d-f) and side (g-i) views for each of the 3 states. The dimer interface helices are colored in green and pink to differentiate individual monomers in the single molecule. Red arrows indicate the distance between the PHY domains, blu e arrows the distance between the top of PCM dimer interface helices and black arrows the distance between the PAS domains. In the Pr state, the dimer interface helices are ∼25 Å apart (a,g). This distance drastically reduces to 14 and 16 Å in the Pr/Pfr heterodimer (b,h) and homodimer Pfr states (c,i), respectively. The PHY domains are pushed apart from ∼22 Å in the Pr state to ∼35 Å in the heterodimer and Pfr states (a -c). Similarly, the PAS domains are pulled in from ∼39 Å in dark state to ∼32 Å in the heterodimer and Pfr states (d -i). The displacement of the domains results in a more parallel arrangement of the dimer interface helices. (e,f). A kink is observed in the coiled coil linkers connecting the HKLD domain to the PCM in Pr/Pfr heterodimer and homodimer in the Pfr state (e,f).

**Figure 5.**
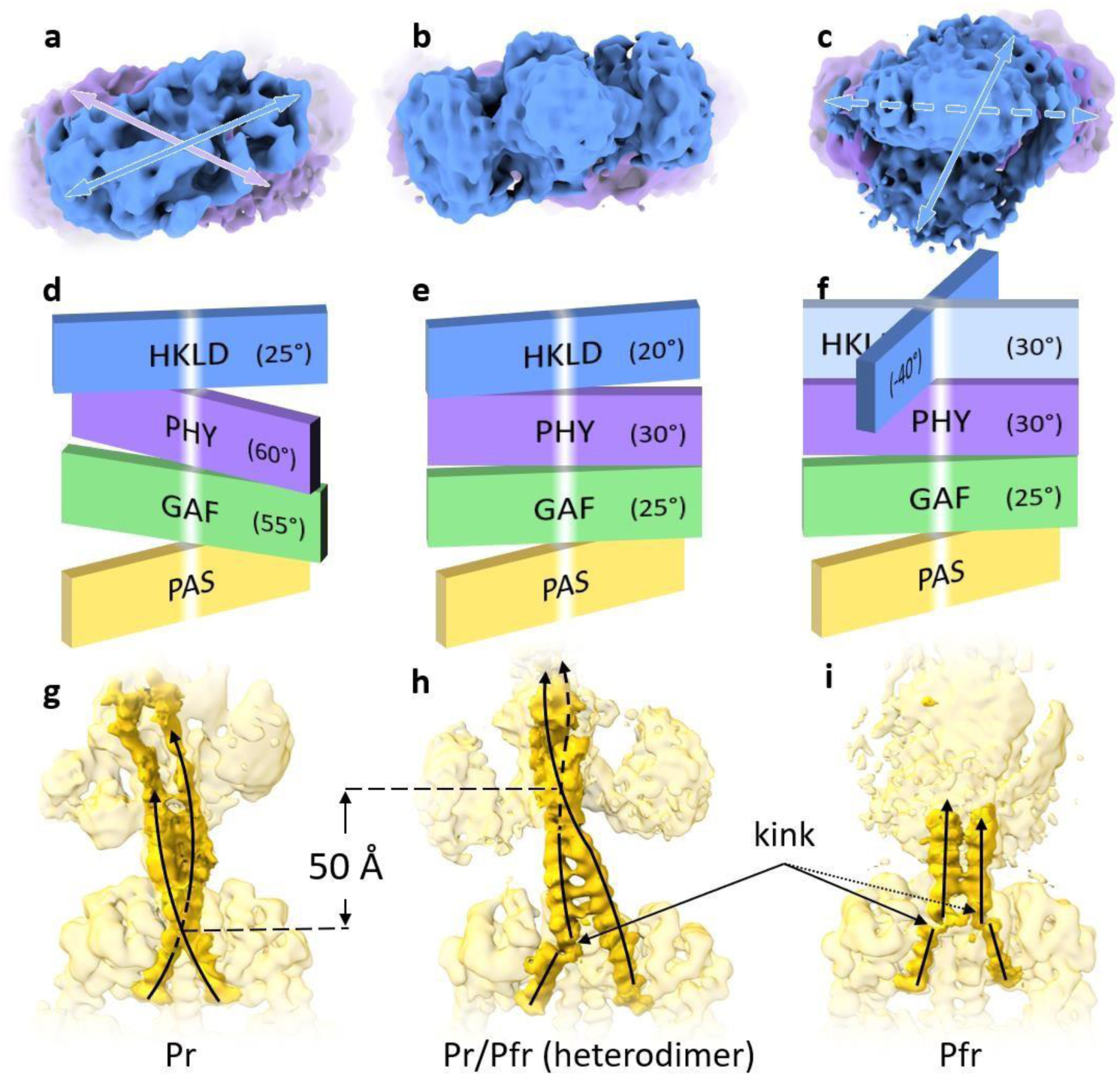
Domain rearrangement s during the Pr/Pfr photocycle. (a _–_ c) Top view of the cryo -EM maps for the HKLD in the Pr homodimer, Pr/Pfr heterodimer and Pfr homodimer. In (a) the arrows indicate the orientation of the PHY (purple) and HKLD (blue) domains. In (c), the solid arrow represents the orientation of the HKLD and the dotted arrow represents an alternate orientation of the HKLD. (d _–_ f) Cartoon represe ntation of the domain orientations for Pr, the hybrid and the Pfr states. The domains are individually colored and marked. The white median represents the dimer interface which separates the two monomers. The orientation of PAS domains in all states are ke pt identical for reference. Clockwise rotation angles are displayed for each domain. The alternate orientation of the HKLD in the Pfr state is shown by an additional light blue panel in (f). (g -i) The orientation of coiled coil linkers (shown in bright yel low) in the three different states. (g) In the Pr state, the linkers cross each other within the PHY domain. (h) The linkers uncoil such that the intersecting point moves 50 Å above towards the HKLD. A kink in the linker helix is observed in subunit A. (i) The linker helices originating from the PCM are parallel to each other. The helical kink is observed in both subunits.

### 2.2 The structure of the Pfr homodimer

The structure of the full length SaBphP2 homodimer in the Pfr state was obtained after irradiating the purified protein with 640 nm light (Fig. 2 c). The two β-strands of the sensory tongue moved apart in both subunits (Fig. 3 d and e). The strand that returns from the BV pocket to the PHY domain has changed to an α-helix. The length of the sensory tongue is now shorter (14 Å compared to 18 Å, Fig. 3 d and e). The BV environment has undergone significant changes. The network of hydrogen bonds formed between the BV and neighboring conserved amino acids that stabilizes the BV in the Pr state is replaced with a new hydrogen bond network (Fig. 3 b) due to significant conformational changes and the sliding of the BV chromophore in the BV-binding pocket of the GAF domain. Ring D no longer forms hydrogen bond with the conserved His275 due to the Z/E isomerization of the C_15_=C_16_ bond and the 180□ rotation of the D-ring (compare Fig. 3 a and b). Notably, the salt bridge of Asp192-Arg457 is broken and the conserved Asp now forms a hydrogen bond with Ser459 of the PRXSF motif as a result of the conformational transition of the sensory tongue from a β-sheet to an □-helix (Fig. 3 b).

Unlike in the Pr state, the dimer interface helices are parallel to each other (Fig. 4 f). The hinge point of the movement is located within the GAF domain (Fig. 4 g,h). The PAS domains are pulled closer by ∼7 Å and the PHY domains are pushed apart by more than 10 Å in the homodimer of the Pfr state. This motion translates to a rotation of the domains. The GAF and PHY domains rotate by ∼30˚ and the HKLD by ∼70˚ compared to the Pr state (Fig. 5e). The coiled linkers straighten up. However, the continuity of the long helix is broken, resulting in a kink in both subunits (arrows in Fig. 4 f, 5 i). In addition to the rotated conformation, the HKLD has an alternate conformation which is oriented at an angle similar to that seen in the Pr state (Fig. 5 c).

### 2.3 The structure of the Pr/Pfr hetrodimer

In addition to the homodimers, a Pr/Pfr heterodimer can be identified in the cryo-EM data. Figure 2 b shows its structure which has been obtained after exposure of the SaBphP2 to ambient white light. Within the heterodimer one monomer is in the Pr and the other in the Pfr state. In the PCM region, the cryoEM map is resolved to better than 3 Å (Supplementary Fig.1 b). Map features in the HKLD region are resolved at ∼9 Å resolution. The map reveals the overall shape of the HKLD (Fig. 2, Supplementary Fig.2) which is consistent with the previously determined crystal structure of a HK (Marina, Waldburger et al. 2005). The backbone features of the protein in the HKLD region become more pronounced after local refinement in *cryoSPARC (Punjani, Rubinstein et al. 2017)* (Supplementary Fig.1 g). The structure of the HK can be modeled and refined in this map (Fig. 2 b, Supplementary Fig.2). The structure of subunit A is similar to that of the Pr state with the BV in the Z-configuration and the sensory tongue in the β-sheet conformation. Similarly, the BV chromophore and the configuration of the sensory tongue in subunit B is essentially identical to that in the Pfr homodimer (Fig. 2). The overall structure of the heterodimer is more similar to that of the Pfr homodimer (Fig. 4 and Fig. 5). The dimer interface helices are parallel like those in the Pfr homodimer and the relative orientations of the PAS, GAF and PHY domains are almost identical (Fig. 5 e). The structures of the coiled coil linker and the HKLD, however, differ. During the unwinding of linker helices the intersecting point is shifted by ∼50 Å towards the HKLD (Fig. 5 g,h). A kink (Fig. 4 e, Fig. 5 h) appears in the linker of subunit A only, which unexpectedly is the monomer still in the Pr state. Despite the structural changes of the PCM, the HKLD itself does not rotate and its orientation remains closer to that in the Pr state (Fig. 5 d,e).

### 2.4 Cryo-EM structures of the PCM

Cryo-EM structures and corresponding maps of the SaBphP2 PCM lacking the OM were captured as Pr/Pfr heterodimer and the Pfr homodimer are shown in Fig. 3 c and f, respectively. The resolution (∼4.5 Å) is lower than that of the corresponding PCM part (∼ 3 Å) in the full-length SaBphP2 structures (Supplementary Table 1) which can be partially ascribed to the smaller size of the PCM (∼120 kDa versus ∼180 kDa of the intact SaBphP2),. The structures of the PCM in two different states are essentially identical to those of the corresponding PCM within the full-length SaBphP2; see Fig. 3 c as an example.

## 3. Discussion

Our results provide detailed structural insight into the myxobacterial two-component signaling pathway, featuring light as an environmental signal in one of the most complex prokaryotic organisms with a unique life cycle and abundant synthesis of secondary metabolites with potential antimicrobial and anticancer properties (Lapuhs, Heinrich et al. 2022). The described cryo-EM structures explain over half a century of structural research on plant and bacterial phytochromes limited by availability of crystals of intact photoreceptors. Based on the comparison of our cryo-EM structures, the following sequence of events characterize the transfer of the light signal to the enzymatic domain. Light absorption causes the BV chromophore in one of the subunits (subunit 1) to undergo Z to E isomerization (Fig. 1 and Fig. 3). The rotation of the BV D-ring causes significant displacement of the residues in the chromophore binding pocket (Fig. 3). The conformational transition of the sensory tongue in subunit 1 is followed by a moderate rotation of the domains (Fig. 2 and Fig. 3). This movement destabilizes the coiled linker in the opposite subunit (subunit 2) that has not isomerized, resulting in a kink in the linker helix (Figs. 2 - 4). Further relaxations of the PCM facilitate the isomerization in subunit 2. This results in the transition of the sensory tongue to a helix and other structural changes (Figs. 4 and 5). A kink in the linker helix of subunit 1 will finally cause the HKLD to rotate (Fig. 5 and 6). The rotation of the HKLD might open or exclude a binding site for the RR. The X-ray structure of an unrelated RR/HK complex (PDB code: 3DGE) shows that the RR interacts mainly with the 4-helix bundle of the HK (Casino, Rubio et al. 2009). The analog region in the SaBphP2 HKLD is shown in Fig. 2 b (dashed box). Any structural change of the 4-helix bundle must have an immediate effect on the binding properties of the RR. Therefore, the rotation observed in the HKLD of SaBphP2 might have a fundamental influence on the binding and dissociation, and consequently on the activation of the RR.

**Figure 6.**
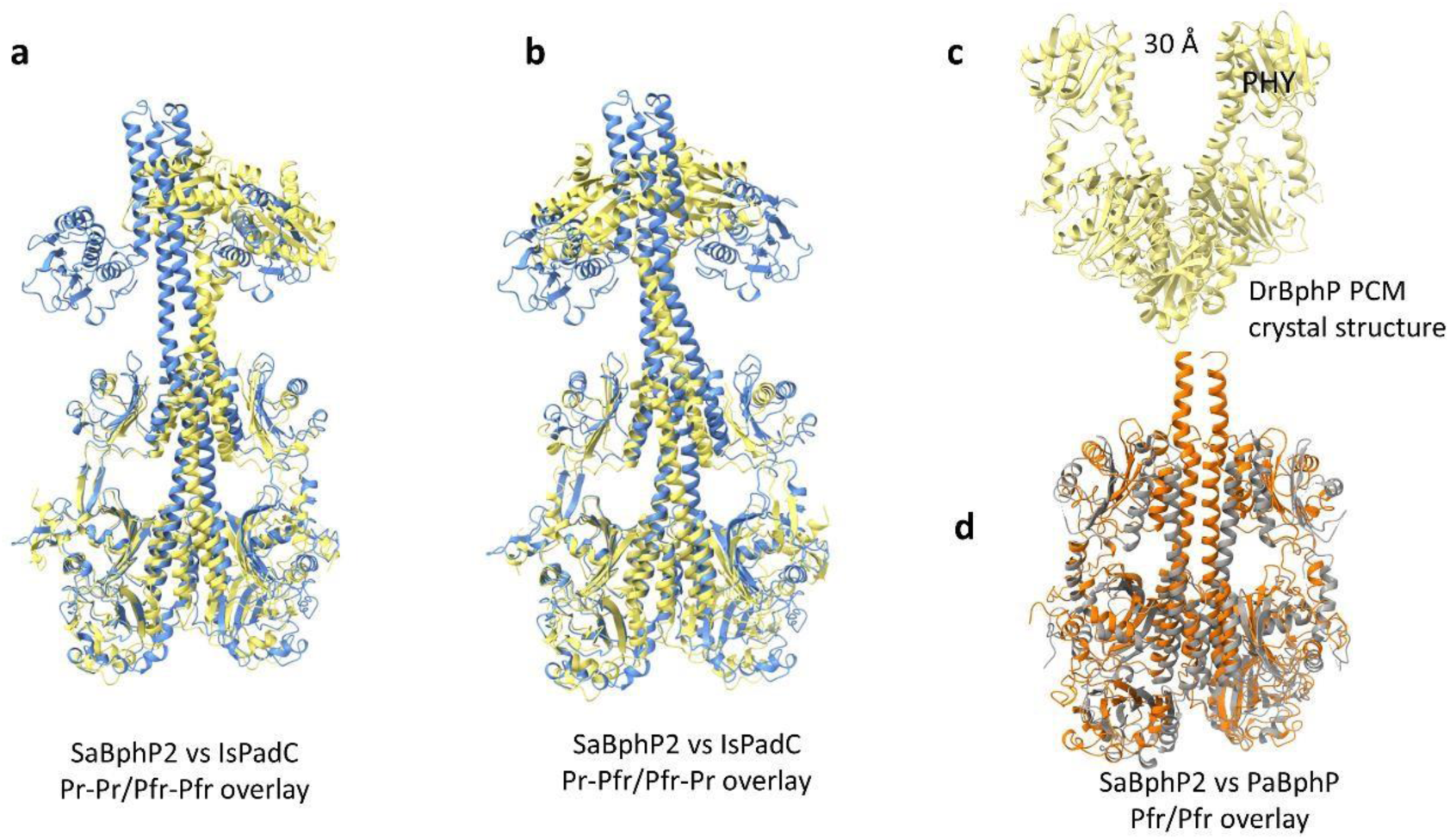
Comparison of intact SaBphP2 with other BphPs. (a -b) Overlay of the Pr/Pfr heterodimer of IsPadC (PDB code: 6ET7, yellow ribbon) on the SaBphP2 heterodimer (blue ribbon). Equivalent monomers in Pr and Pfr of each protein were compared, respectively. The ov erlay is marked Pr -Pr/Pfr-Pfr. The IsPadC heterodimer also features a bent coiled coil linker which is identical to the bend in the SaBphP2 structure, but shifts the OM in the opposite direction. (b) When overlaying the Pr with the Pfr subunits, a bett er alignment of the linker region is achieved. The overlay is marked Pr -Pfr/Pfr-Pr. (c) Crystal structure of the DrBphP PCM in the Pfr state (PDB code: 4O01). Unlike PaBphP and SaBphP2, the DrBphP in the Pfr state shows a large opening of the PHY domain. (d) Comparison of SaBphP2 in Pfr (orange) state with the bathy phytochrome PaBphP (PDB code: 3G6O, gray ribbon) in the Pfr state. The dimer helices are parallel in both proteins. In PaBphP monomers are separated further than in SaBphP2.

In the related classical phytochrome DrBphP, it was shown that the dimer interfaces adjacent to PHY domains separate by up to ∼30 Å (Fig. 6 c) upon photoconversion (Takala, Bjorling et al. 2014, Burgie, Zhang et al. 2016). More recently, it was concluded with cryo-EM studies that there is no significant change in the position of the PHY domain upon photoconversion of DrBphP (Wahlgren, Claesson et al. 2022). Our observations contradict both conclusions. We observe a ∼14 Å displacement of the PHY domains (Fig. 4 a,b) but in a manner quite different from previous observations. The published cryo-EM structure (Wahlgren, Claesson et al. 2022) is determined from a genetically modified photochrome construct where a RR protein that is about the size of the OM itself has been covalently linked to the phytochrome. The heavy addition might hinder the necessary large-scale rotations of the OM. On the other hand, the crystal structures in the Pfr states were from truncated DrBphP without the OM. The very large separation of the PHY domains may have been introduced during crystal formation and then exacerbated by the absence of constraints that prevent the PHY domain from moving beyond the physiological range. Moreover, the cryoEM structures of SaBphP2 in the Pfr state (in the full length and truncated forms) are more in line with the crystal structure of the bathy phytochrome of *P. aeruginosa* (Yang, Kuk et al. 2008) (Fig. 6 d) which in the Pfr state does not show large PHY domain separation and features a parallel dimer interface.

The SaBphP2 investigated here is a wild type protein, containing both PCM and OM. As such, the structure is free of restraints that prevent natural motions (including steric and packing clashes present in crystals), and the freedom that may result from truncation. Burgie and coworkers also made a similar observation to ours with full length DrBphP using cryo-EM (Burgie, Bussell et al. 2014). Despite low (∼27 Å) resolution, a ∼35˚ rotation and a ∼9 Å movement of the PHY domains has been determined (compared to ∼30˚ and ∼13 Å seen here) without the large PHY domain opening determined from crystal structures. In addition, X-ray solution scattering studies (Bjorling, Berntsson et al. 2016, Takala, Niebling et al. 2016) and NMR investigations support our findings (Isaksson, Gustavsson et al. 2020).

Both, the PCM and the HKLD are large moieties. A linker that consists of a single α-helix connects both modules. As a result, the dimer interface beyond the PHY domain is formed by a flexible coiled coil (Fig. 2 and Fig. 5) that propagates the signal. When the PCM is photoexcited the HKLD cannot react instantaneously. A kink is formed in one of the linker helices when the rotation of one module lags behind the other (Fig. 5 h). The kink forms near Phe485. Aromatic residues, especially phenylalanine, are known to have high propensity for helical disruption (Huang and Chen 2012, Wilman, Shi et al. 2014). Several studies have also reported asymmetric kinking in the interface helices in other HK structures including a BphP from a plant pathogen (Otero, Klinke et al. 2016) (Fig. 6 a,b). They were associated with signal transduction and modulation (Diensthuber, Bommer et al. 2013, Ferris, Coles et al. 2014). Since the kink appears both in crystal and cryo-EM structures, it seems physiologically relevant. It has been proposed that the change in the hydration environment caused by the exposure of the Phe485 hydrophobic sidechain to the solvent breaks the hydrogen bonds that are stabilizing the helix (Otero, Klinke et al. 2016). However, our experiments favor the view that the helix is destabilized by the β-sheet to α-helix transition of the sensory tongue in the opposite subunit that causes substantial structural relaxations along the dimer interface (Fig. 5 h). It seems that the formation of the second kink is the decisive structural reason for the subsequent rotation of the OM.

Most crystal structures of BphPs are truncated homodimers lacking the OM domain (Takala, Bjorling et al. 2014, Burgie, Zhang et al. 2016, Sanchez, Carrillo et al. 2019, Carrillo, Pandey et al. 2021). Crystals tend to be selective for proteins with the same structure. This could have resulted in an apparent homogeneity in crystals of the SaBphP2 resting state. However, the Pr/Pfr heterodimer (Fig. 2 b) is most likely dominant in solution and is probably prevalent *in vivo*. In plants, heterodimerization is integral for signaling and the phytochrome can exist as a heterodimer in the resting state (Sharrock and Clack 2004). In contrast, the existence of a full-length BphP heterodimer has not been expected, although heterodimeric structures were characterized with modified BphPs that have a different enzymatic domain (diguanylyl cyclase) (Gourinchas, Heintz et al. 2018) (Fig. 6 a,b). The hybrid structure presented here shines light on the significance of the heterodimers. Comparing the structures between Pr, hybrid and Pfr states, it appears that the Pr/Pfr heterodimer is, in some sense, an intermediate in the photocycle. When the BphP exists as a heterodimer in its resting state, the conversion of a heterodimeric form to the Pfr state is expected to be faster and more efficient than the direct conversion which has been already established in plant phytochromes (Klose, Venezia et al. 2015).

Since the BphPs are part of the important two-component signaling mechanisms, it would be desirable to determine the structure of the BphP in complex with a RR. This will clarify where and to which of the states the RR binds. The conformational change of the sensory tongue and the twisting of the PCM dimer interface helices are coupled. Signal propagation via both the sensory tongue and the dimer interfaces is suggested but the decisive conclusion is missing. Time-resolved (TR) cryo-EM presents a unique opportunity to study these large-scale conformational changes. Initial successes with TR cryo-EM are reported (Frank 2017, Dandey, Budell et al. 2020, Torino, Dhurandhar et al. 2023). The TR cryo-EM experiment will reveal the specific order of conformational changes that ultimately trigger the enzymatic activity and whether the Pr/Pfr heterodimer is an important photocycle intermediate and/or part of the innate ground state heterogeneity.

## 4. Material and Methods

### Protein purification

The coding region of the wild-type SaBphP2 were PCR-amplified from *S. aurantiaca* DW4/3–1 genomic DNA and cut by restriction enzymes NdeI and HindIII (New England Biolabs, Beverly, USA), and ligated into the corresponding sites of the expression vector pET28c(+) (Invitrogen, Carlsbad, CA). The constructed plasmids and the pET11a vector carrying heme oxygenase were transformed into *Escherichia coli* BL21 (DE3) strain for expression. Cells were grown at 37° C to a OD600 value of 0.6 followed by induction with 1 mM IPTG and addition of 0.5 mM δ-aminolaevulinic acid (DAC) overnight. Cells were recovered in 150 mM NaCl, 20 mM Tris-HCl, pH 8.0 and 15% v/v glycerol with protease inhibitor. Lysis was performed with pulse sonication on ice bath. The insoluble cell debris was removed by highspeed centrifugation, and the supernatant was incubated with 200 μM BV in Dimethyl sulfoxide (DMSO) in the ratio 1:100 for 30 min at 4°C. The solution was applied to Talon Co^+2^ metal ion affinity chromatography column. The column was washed with high salt buffer (20 mM Tris-HCl, 1 M NaCl, pH 8.0) followed by low salt buffer (20 mM Tris-HCl, 1 M NaCl, pH 8.0) with 20 column volumes each. The protein was eluted by 300 mM imidazole, 20 mM NaCl, 20 mM Tris-HCl, pH 8.0. After elution the protein was immediately transferred into stabilizing buffer containing 20 mM NaCl, 20 mM Tris-HCl, pH 8.0. All steps were performed under green safety light. The purified protein was immediately frozen at -80° C and shipped on dry ice to the Simons Electron Microscopy Center (SEMC) of the New York Structural Biology Center (NYSBC).

### Grid preparation

UltrAuFoil holey grids (300 mesh R1.2/1.3, Quantifoil) were plasma-cleaned for 7 s using oxygen and argon gases with a Solarus II Gatan Plasma System. The grids were vitrified by plunging into liquid ethane on an FEI Vitrobot automatic plunge freezer with a blotting time of 1 seconds and incubation time of 10 seconds. To obtain the full-length SaBphP2 Pr/Pfr hybrid, the protein was frozen at a concentration of 0.3 mg/mL with a blotting time of 1.5 seconds under standard laboratory fluorescent light conditions. To obtain SaBphP2 in the all-Pr and all-Pfr states, sample grids were prepared under green safety light, at a concentration of 0.4 mg/mL. The all-Pr state was produced by pre-illumination with 740 nm light, while the all-Pfr state was obtained by pre-illumination with 640 nm light. Grids with the truncated PCM construct were prepared under green safety light, at a concentration of 0.7mg/mL and pre-illuminated with 640 nm light to obtain the all-Pfr state. To obtain the PCM construct in the hybrid state the grid was not pre-illuminated, and frozen under standard laboratory fluorescent light conditions. All frozen grids were clipped in AutoGrid cartridges (Thermo Fisher Scientific) and stored in liquid nitrogen until data acquisition.

### Data Acquisition

All grids were imaged on a Thermo Fisher Scientific Titan Krios microscope equipped a Gatan BioQuantum K3 energy filter direct electron detector camera. Movies with SaBphP2 in the all-Pr state were collected in counting mode with a 2000 ms exposure time, 50 frames, 40 ms per frame and a total dose of 56.39 e-/Å2. A total of 15,533 movies were collected at a pixel size of 0.844 Å/px. Movies of SaBphP2 in the all-Pfr state were collected in counting mode with a 2000 ms exposure time, 50 frames, 40 ms per frame and a total dose of 59.31 e-/Å2. A total of 19,607 movies were collected at a pixel size of 0.844 Å/px. Movies of the SaBphP2 Pr/Pfr hybrid were collected in counting mode with a 2000 ms exposure time, 50 frames, 40 ms per frame and a total dose of 67.02 e-/Å2. A total of 17,880 movies were collected at a pixel size of 0.844 Å/px. Movies on the (truncated) PCM in the all-Pfr state were collected in counting mode with a 2000 ms exposure time, 50 frames, 40 ms per frame and a total dose of 49.78 e-/Å2. A total of 19,607 movies were collected at a pixel size of 0.844 Å/px. Movies on the PCM in the hybrid state were collected with a 2000 ms exposure time, 50 frames, 40ms per frame and a total dose of 49.78 e-/Å2. A total of 17,707 movies were collected with a super-resolution pixel size of 0.422 Å/px. All data acquisition was done using Leginon (Suloway et al., 2005).

### Cryo-EM map reconstruction and structure determination

Data were processed for all structures in a similar manner starting with the Pr/Pfr heterodimer. Data processing was done using *CryoSPARC* (v4.2.1) (Punjani et al., 2017). The processing and refinement statistics are shown in Supplementary Table 1.

### SaBphP2 cryo-EM structures

The full-length SaBphP2 structure of the homodimer in the Pr state has been determined at an overall resolution of 4.13Å. The structure of the homodimer in the Pfr state has been determined at 3.75Å resolution. The Pr and Pfr structures were obtained by illuminating the protein with the far-red and red light respectively. Interestingly, when BphP is handled under ambient white light, a heterodimer structure, with one monomer in Pr and the other in the Pfr state is identified. The cryo-EM map of the Pr/Pfr heterodimer is reconstructed at 3.75Å overall resolution. The local resolution of the PCM in all three full-length SaBphP2 structures is around or better than 3Å (fig. S1). The cryo-EM maps for a truncated SaBphP2 containing the PCM only were obtained at a resolution of ∼4.5 Å (Supplementary Table 1). Structures of a homodimer in the Pfr state and the one of a heterodimer with individual monomers in distinct Pr and Pfr states in the same molecule, respectively, were determined.

### The Pr/Pfr heterodimer of the full-length SaBphP2

The 17,880 raw movies were pre-processed with *patch motion correction* and *patch CTF estimation* jobs. The resulting micrographs were subjected to exposure curation from which 13,321 exposures were selected. The rest with poor CTF fits and large full-frame motions were eliminated. *Blob picking* was used to pick the initial lot of particles. Using 68,220 particles identified as full length phytochrome in 2D classification job, a low-resolution map at ∼10 Å was generated with *ab-initio reconstruction*. The resulting map was used for template-based particle picking. This yielded approximately 270 particles per micrograph. Following an inspection of the particle picks, a total of 1,520,224 particles (box size 400 x 400 pixels) were extracted and averaged using the *2D classification* utility. High resolution classes were selected and subjected to further rounds of *2D classification.* Several rounds of particle curation resulted in a dataset with 865,248 particles. These particles were used by an *abinitio reconstruction* job to generate five 3D maps without any reference. The five *ab-initio* classes were refined and the particles were classified amongst them with *heterogenous refinement*. The individual maps were refined to high-resolution and validated using the gold standard FSC with *nonuniform refinement*. A final map with best FSC resolution of 3.75 Å (Supplementary Fig.2) was obtained with 215,374 particles.

The crystal structure of the PCM in the Pr state fits reasonably well within subunit A of the cryo-EM map (Fig. 2 a). The structure could be refined with minor adjustments. Subunit B, however, was in the Pfr state and the crystal structure did not fit. The Pfr structure determined here is quite different from the structures of other BphPs in the Pfr state. Several sections were identified in the Pr state where the secondary structures were conserved in the Pfr state as well. These sections were isolated from the crystal structure and fitted individually to the cryo-EM map sections. The remaining residues were manually built with *Coot* (Emsley, Lohkamp et al. 2010) and *ChimeraX* (Pettersen, Goddard et al. 2021) as the side chains were clearly visible in the sharpened map produced by the *nonuniform refinement* job in *CryoSPARC*. The linker region was also modeled this way. Once an approximate model was obtained, real-space refinement in *phenix* (Liebschner, Afonine et al. 2019) was used to refine the model against the unsharpened map. To model the HKLD at a resolution of ∼9 Å, its structure was predicted by *Alphafold (Jumper, Evans et al. 2021)*. The cryo-EM map in the HKLD region is hemispherical (Supplementary Fig.2). The HKLD is connected to the dimer helices with a single stranded loop. If the catalytic domain was rotated by 90° relative to the *AlphaFold* solution, it would still fit into the overall cryo-EM map envelope. The two possible orientations can be distinguished by the correlation factor between the structure and the cryo-EM map that becomes available after refinement. The model was fitted into the map and merged with the previously refined structure of the PCM and the linker. The overall structure was refined with secondary structure restraints turned on. The correlation factor is 53 % for the *AlphaFold* solution. With a correlation factor of 65 % the rotated orientation is favored (Fig. 2, Supplementary Fig.2 e).

### The Pr homodimer of the full-length SaBphP2

Processing steps for the Pr dataset follow similar steps as for the hybrid dataset except this time, the hybrid map was available and used as the template for particle picking. A total of 1,112,759 particles were picked. These particles were filtered through multiple runs of *2D classification*. A set of 726,193 particles were used to reconstruct five *ab initio* maps. The best three maps were used as initial maps for *heterogenous refinement*. The map with best resolved features were used for further *homogenous* and non-*uniform refinement*. The final map with GSFSC resolution of 4.13 Å was obtained from 377,549 particles (Supplementary Fig.3). The PCM of subunit A of the hybrid structure was used to model the the PCM on both subunits in Pr state. The coiled coil linker was manually modeled in *Coot*. The overall structure was refined in *phenix*.

### The Pfr homodimer of the full-length SaBphP2

The Pfr dataset processing pipeline followed the same procedure as above. Starting with 2,217,983 particles picked with template, a final map with GSFSC resolution of 3.75 Å (Supplementary Fig.4) was obtained from 155,342 particles. The PCM of subunit B of hybrid structure was used to model the PCM on both subunits in Pfr state. The coiled coil linker was manually modeled in *Coot* and refined in *phenix*.

### The Pr/Pfr heterodimer of the truncated SaBphP2 PCM

Since the dataset for the PCM heterodimer was collected in super resolution mode, an *output F-crop factor* of 1/2 was applied during *Patch motion correction* job in *CryoSPARC*. A template was created using the PCM portion of the map obtained from the full-length heterodimer. A total of 2,903,962 particles were extracted using a smaller box size (compared to full-length dataset) of 256 x 256 pixels. The particles were filtered through multiple rounds of 2D classification (Supplementary Fig.5 a). The final map with GSFLC resolution of 4.5 Å was obtained from 301,363 particles. The PCM structure extracted from the full-length heterodimer was used as an initial model for refinement in *Phenix*. The chromophore was omitted from the model prior to refinement.

### The Pfr homodimer of the truncated SaBphP2 PCM

The dataset of the truncated SaBphP2 was processed (Supplementary Fig.5 b) in the same way as the dataset of the respective full-length construct, since in both cases the movies were collected without the super resolution mode. The only difference was the particle extraction size which was set to 256 x 256 pixels. Starting with 3,616,004 particles, a final map was reconstructed from 276,639 particles at a GSFSC resolution of 4.61 Å. The PCM part of the full-length SaBphP2 Pfr homodimer was used as an initial model for refinement in *phenix*. The BV chromophore was omitted from the model prior to refinement.

## Data and materials availability

All relevant data are included with the paper and/or are available from the corresponding authors upon reasonable request. Cryo-EM maps and atomic models were deposited to the Electron Microscopy Data Bank (EMDB) and Protein Data Bank (PDB) databases. The PDB codes are: 8UPH (full-length Pr state), 8UPM (full-length Pfr state), 8UPK (full-length hybrid state), 8UQK (PCM Pfr state) and 8UQI (PCM hybrid state). The EMDB accession codes are EMD-42448 (full-length Pr state), EMD-42452 (full-length Pfr state), EMD-42450 (full-length hybrid state), EMD-42469 (PCM Pfr state) and EMD-42472 (PCM hybrid state). Atomic coordinates of other phytochromes used for comparison in this study are available in the PDB under accession codes 6ET7, 3G6O and 4O01 respectively. All data are available in the main text or the supplementary material.

## 6. Supplementary Materials

**Supplementary Figure 1.**
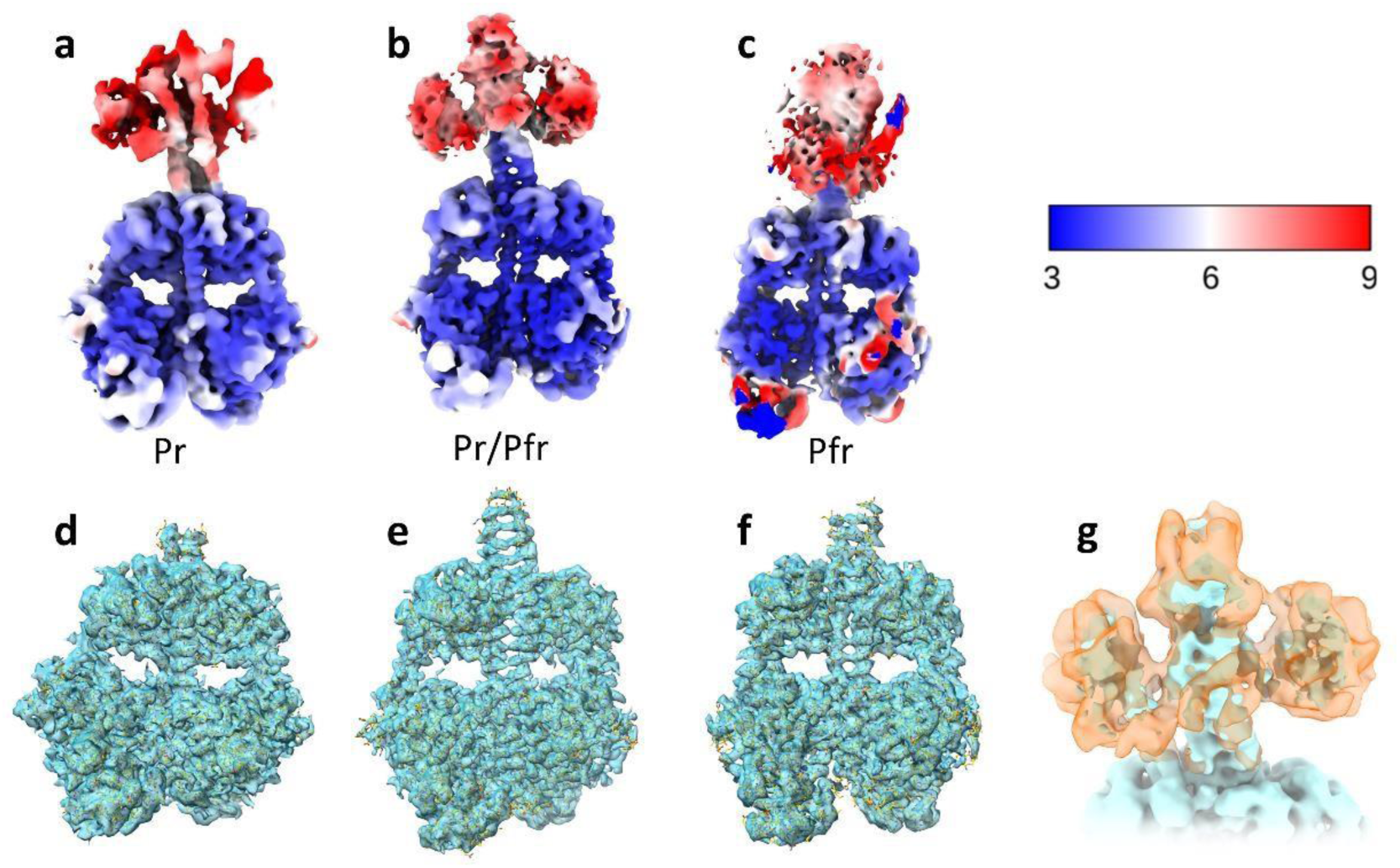
Differences in local resolution of the reconstructed full length BphP maps. (a-c) The local resolution map of the full length phytochrome in Pr, Pr/Pfr heterodimer and Pfr states. The numbers in the color key are in Ångstroms. (d-f) The sharpened map of the PCM in the full-length BphP in the Pr, Pr/Pfr and Pfr states. The maps are shown in transparent teal surface and the atoms and bonds are shown in orange sticks. The side chains are clearly distinguishable in the maps into which the atomic models can be fit perfectly. (g) The map features of the HKLD region in the Pr/Pfr heterodimer map improve after focus refinement (transparent orange) compared to before (blue). The protein backbone can be traced but the side chains are not resolved.

**Supplementary Figure 2.**
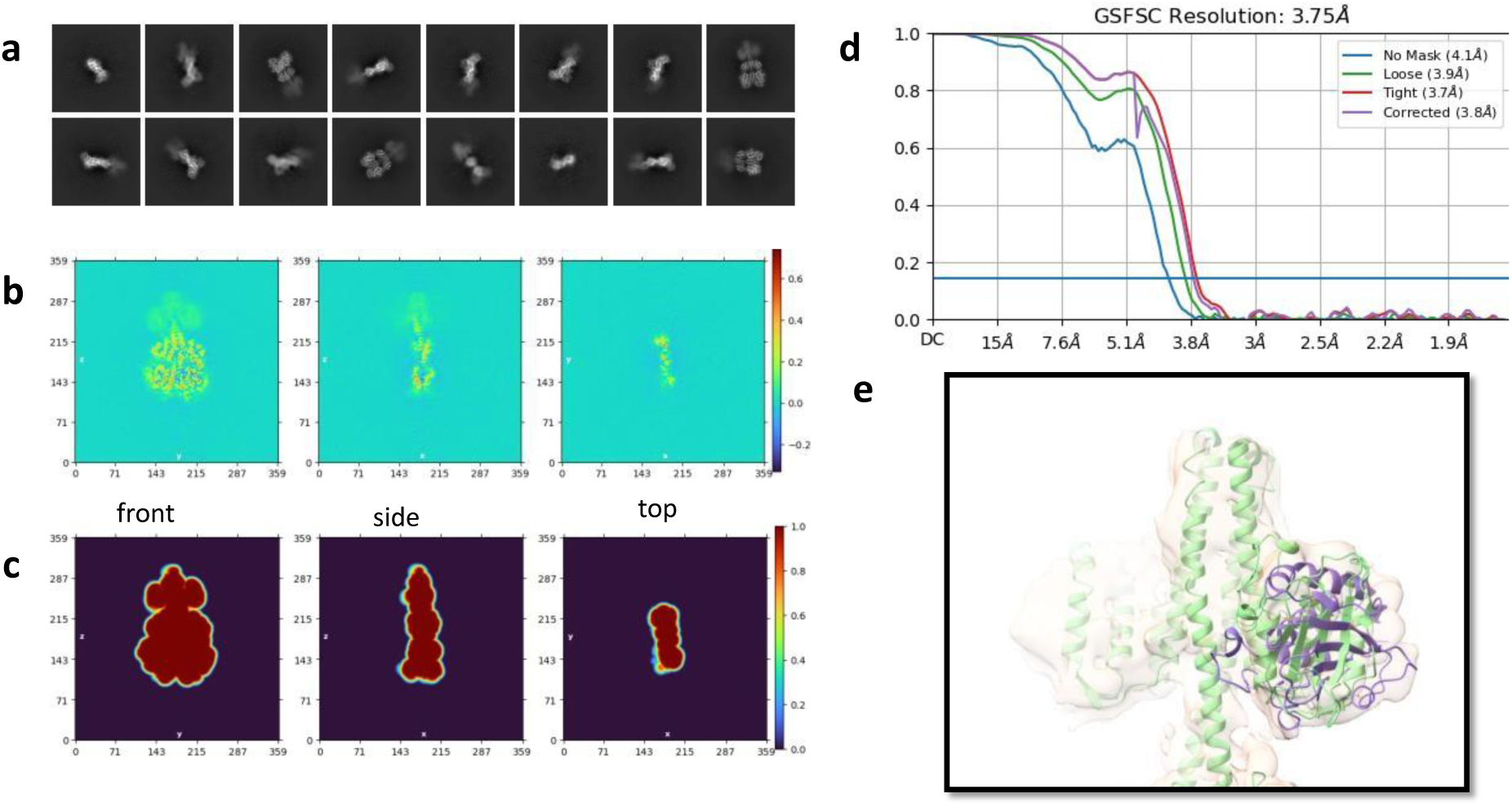
Reconstruction of the cryo-EM map of the full-length SaBphP2 Pr/Pfr heterodimer. (a) 2D class averages of the particles in hybrid state. (b) Real space slice of the map during the final iteration of refinement. The shape of the HKLD is visible. The left and right side of the map are asymmetric. The α-helix in the sensory tongue is distinguishable on the right side of the map. (c) A real space slice through of the mask used for map refinement. (d) The GSFSC of the reconstructed map is plotted against resolution for several reconstruction methods. (e) The modeling of the HKLD structure into the cryo-EM map. The catalytic domain can be fitted in two different orientations. The original *Alphafold* predicted structure is shown in green cartoon, and the alternate configuration in which the domain is rotated 90 anti-clockwise is shown in purple ribbon. The correlation coefficient for alternate configuration is higher than that for the *Alphafold* prediction.

**Supplementary Figure 3.**
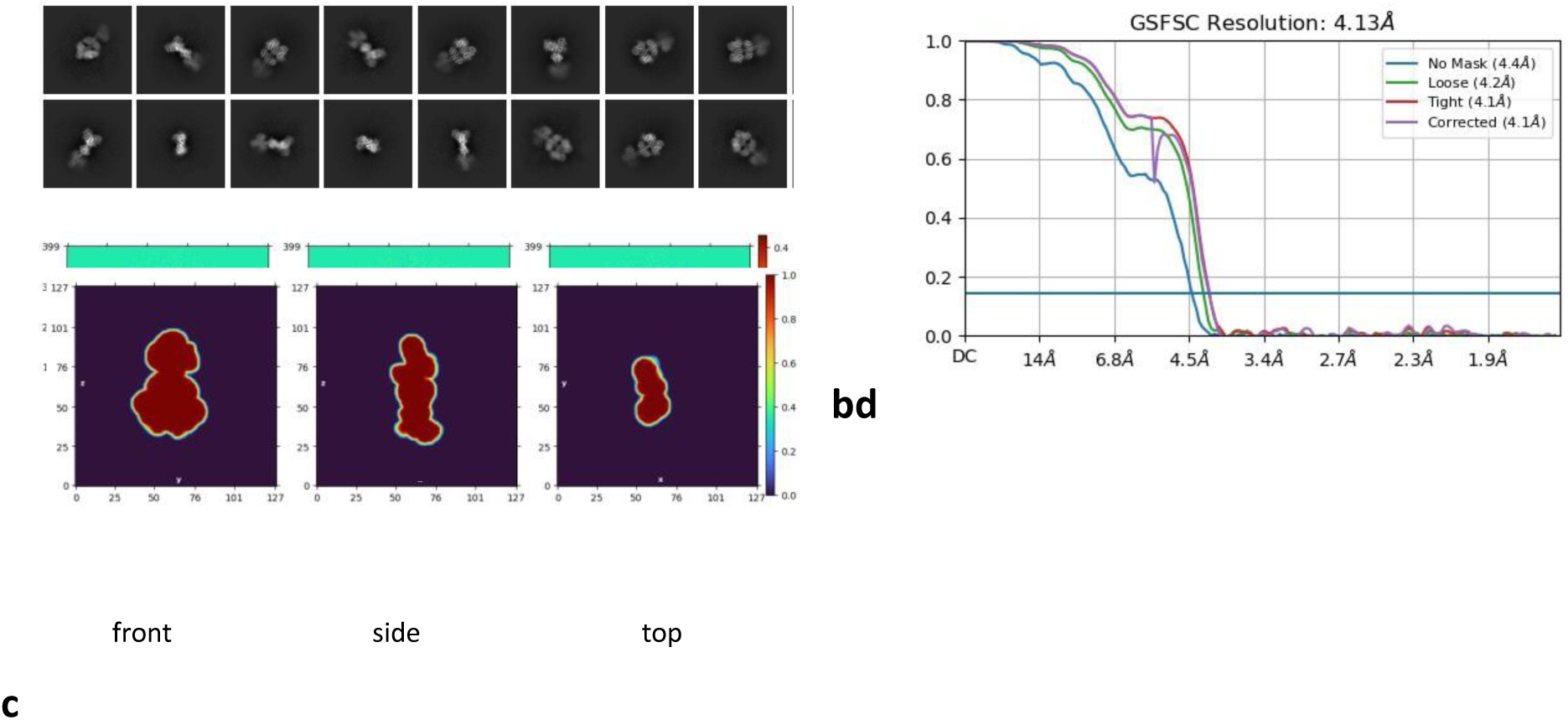
The reconstruction of the cryo-EM map of the full-length SaBphP2 in the Pr state. (a) Examples of 2D class averages of the particles. (b) Real space slice of the map during the final iteration of refinement. (c) A real space slice through the mask used for map refinement. (d) The gold standard Fourier shell correlation (GSFSC) of the reconstructed cryo-EM map is plotted against resolution for several reconstruction methods.

**Supplementary Figure 4.**
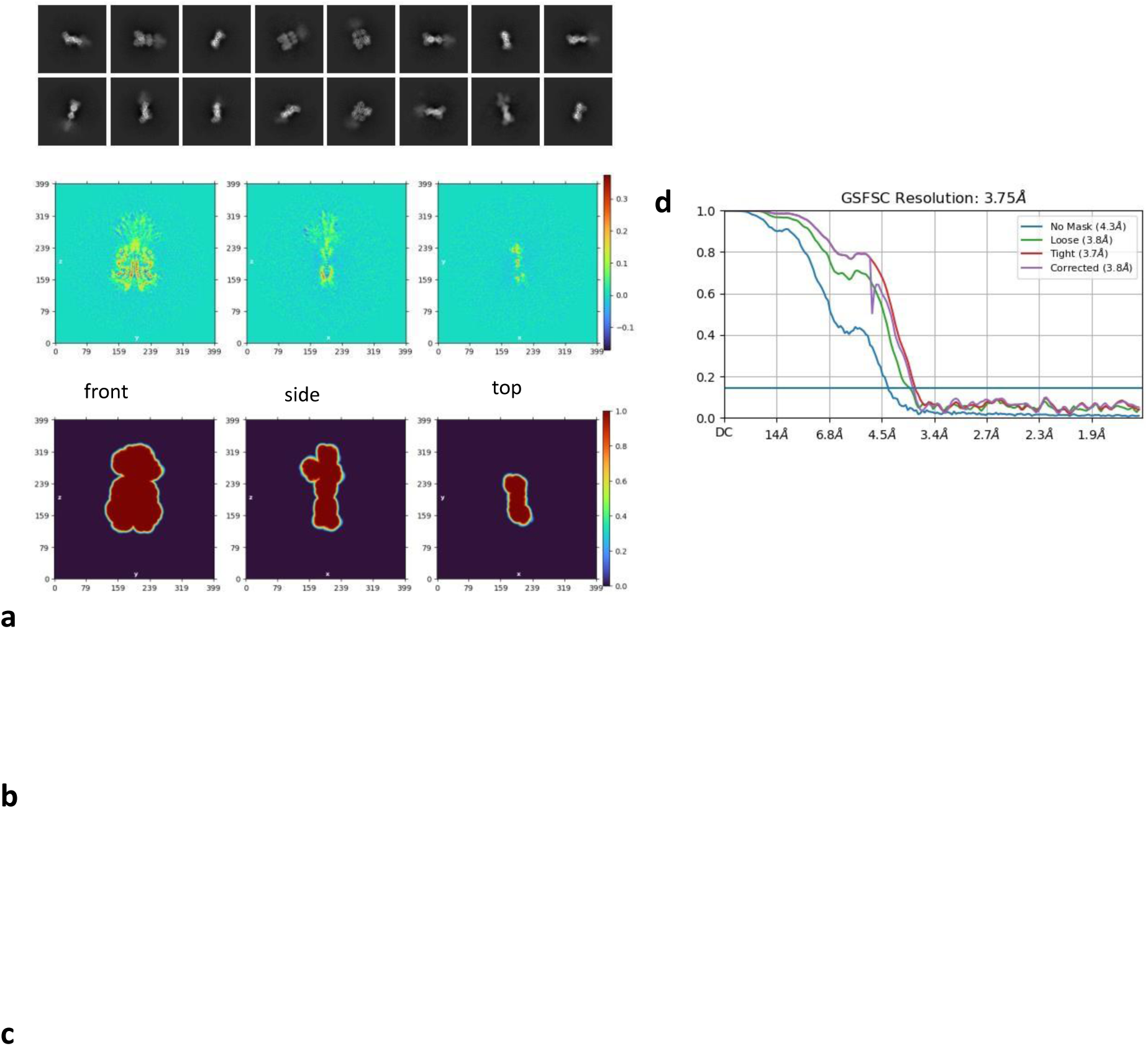
Reconstruction of the cryo-EM map of the full-length Pfr state. (a) Examples of 2D class averages. (b) Real space slice of the map during the final iteration of refinement. The helical sensory tongues are distinguishable. (c) A real space slice through of the mask used for map refinement. (d) The GSFSC of the reconstructed map is plotted against resolution for several reconstruction methods.

**Supplementary Figure 5.**
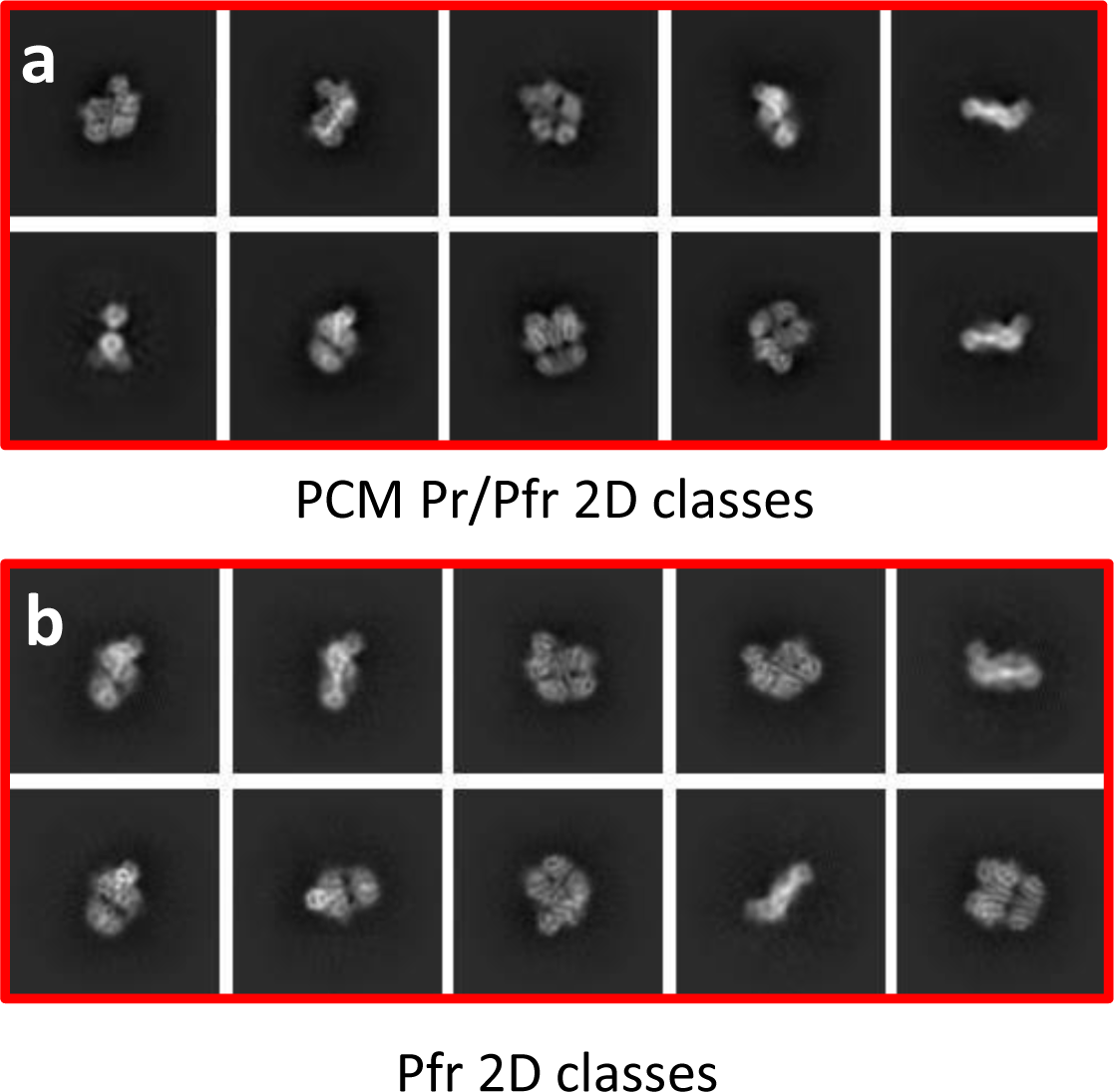
2D class averages of the PCM particles (a) in the Pr/Pfr heterodimeric state, and (b) from Pfr homodimers.

**Supplementary Table 1.**
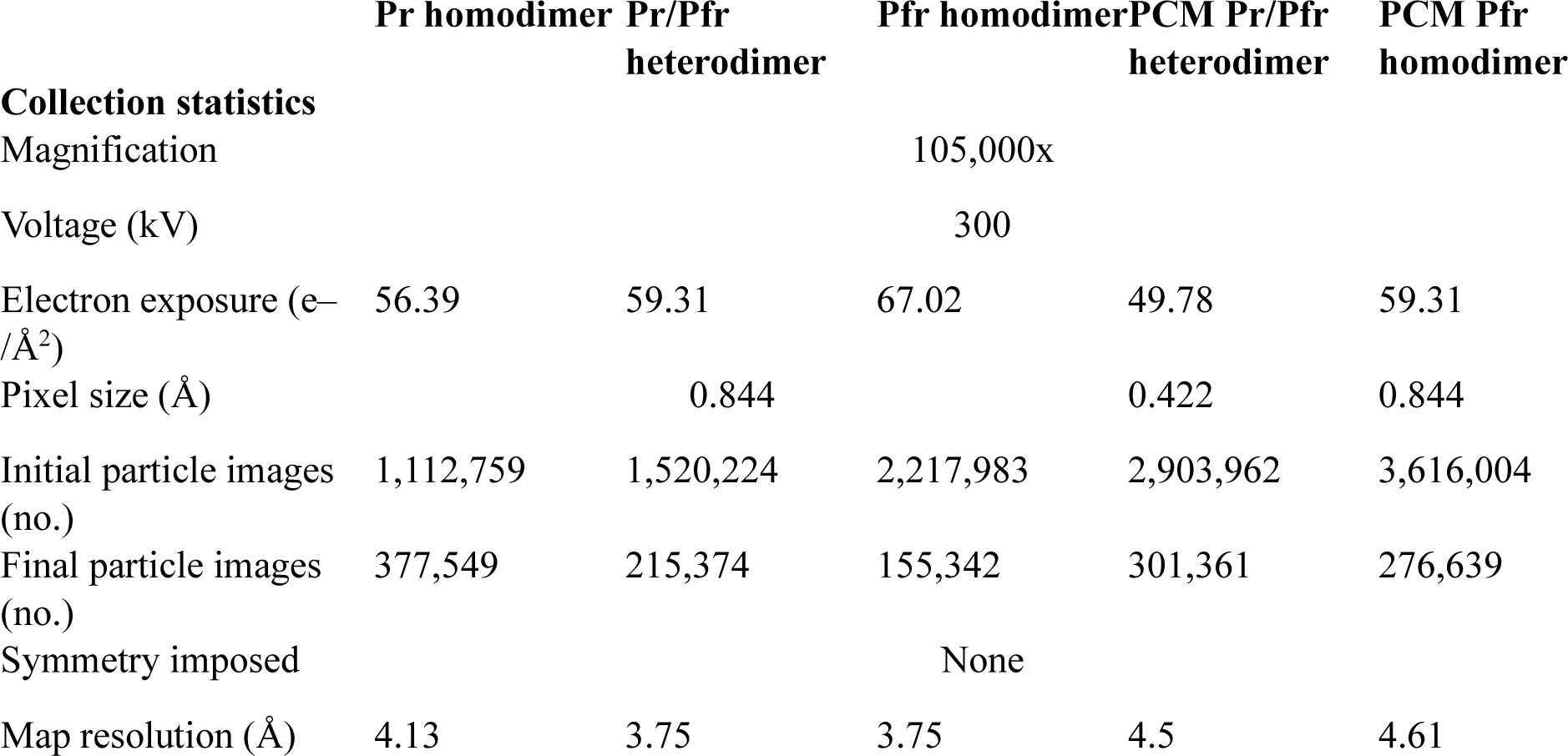
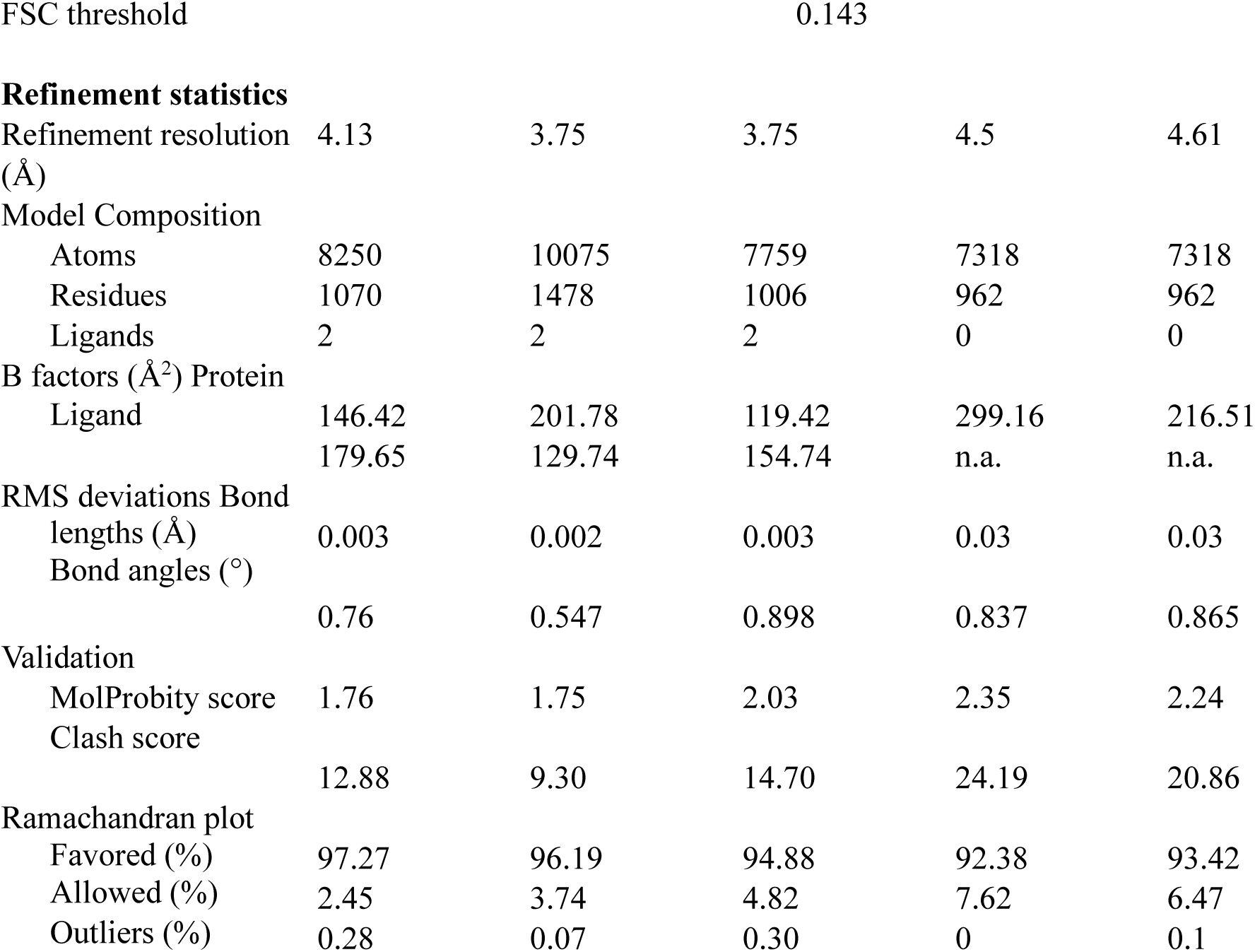
Data collection and refinement statistics.

